# CTCF and Cohesin Regulate Chromatin Loop Stability with Distinct Dynamics

**DOI:** 10.1101/093476

**Authors:** Anders S. Hansen, Iryna Pustova, Claudia Cattoglio, Robert Tjian, Xavier Darzacq

## Abstract

Folding of mammalian genomes into spatial domains is critical for gene regulation. CTCF and cohesin control domain location by folding domains into loop structures, which are thought to be highly stable. Combining genomic, biochemical and single-molecule imaging approaches, we show that although CTCF and cohesin can physically interact, CTCF binds chromatin much more dynamically than cohesin (~1 min *vs.* ~22 min residence time). Moreover, after unbinding, CTCF quickly rebinds another cognate site unlike cohesin (~1 min *vs.* ~33 min). Thus, CTCF and cohesin form a rapidly exchanging “dynamic complex” rather than a typical stable complex. Since CTCF and cohesin are required for loop domain formation, our results suggest that chromatin loops constantly break and reform throughout the cell cycle.

## Introduction

Mammalian interphase genomes are functionally compartmentalized into Topologically Associating Domains (TADs) spanning hundreds of kilobases. TADs are defined by frequent chromatin interactions within themselves while insulated from adjacent TADs (Dekker and Mirny, 2016; Dixon et al., 2012; Hu et al., 2015; Merkenschlager and Nora, 2016; Nora et al., 2012; Wang et al., 2016). Most TAD or domain boundaries are strongly enriched for CTCF (Ghirlando and Felsenfeld, 2016), an 11-zinc finger DNA binding protein (Figure 1A), and cohesin (Skibbens, 2016), a ring-shaped multi-protein complex composed of Smc1, Smc3, Rad21 and SA1/2 that is thought to topologically entrap DNA (Figure 1B). This subset of TADs, which tend to be demarcated by convergent CTCF binding sites, are referred to as loop domains (Rao et al., 2014). Targeted deletions of CTCF binding sites demonstrate that CTCF causally determines loop domain boundaries (Sanborn et al., 2015; de Wit et al., 2015). Moreover, disruption of loop domain boundaries by deletion or silencing of CTCF binding sites allows abnormal contact between previously separated enhancers and promoters, which can induce aberrant gene activation leading to cancer (Flavahan et al., 2015; Hnisz et al., 2016a) or developmental defects (Lupianez et al., 2015). Yet, despite much progress in characterizing TADs and loop domains, how they are formed and maintained remains unclear. Since CTCF and cohesin control domain organization, here we investigated their dynamics and nuclear organization using single-molecule imaging in live cells.

**Figure 1.**
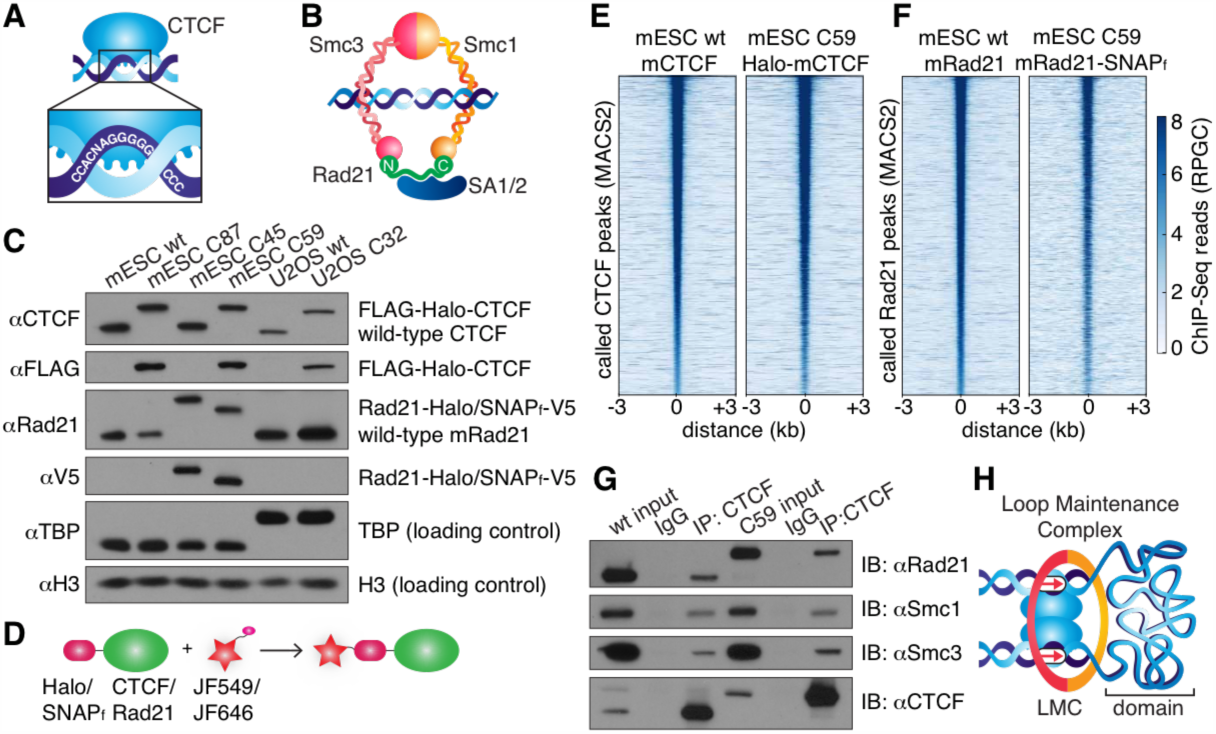
CTCF and Cohesin can be endogenously tagged and form a complex. **(A)** Sketch of CTCF and its DNA binding site. **(B)** Sketch of cohesin, with subunits labeled, topologically entrapping DNA. **(C)** Western blot of mESC and U2OS wild-type and knock-in cell lines. **(D)** Sketch of covalent dye-conjugation for Halo or SNAP_f_-Tag. **(E)** CTCF ChIP-Seq read count, (Reads Per Genomic Content) for wild-type and C59 plotted at called wt-CTCF peak regions. **(F)** Rad21 ChIP-Seq read count, (Reads Per Genomic Content) for wild-type and C59 plotted at called wt-Rad21 peak regions. **(G)** Co-IP. CTCF was immunoprecipitated and we immunoblotted for cohesin subunits Rad21, Smc1 and Smc3. **(H)** Sketch of a Loop Maintenance Complex (LMC) composed of CTCF and cohesin holding together a spatial domain.

## Results

In order to image CTCF and cohesin without altering their endogenous expression levels, we used CRISPR/Cas9-mediated genome editing to homozygously tag mCTCF and mRad21 with HaloTag in mouse embryonic stem (mES) cells (Figure 1C, clones C87 and C45). We also generated a double Halo-mCTCF/mRad21-SNAP_f_ knock-in mESC line (Figure 1C, C59) as well as a Halo-hCTCF knock-in human U2OS cell line (Figure 1C, C32). Halo- and SNAP_f−_Tags can be covalently conjugated with bright cell-permeable small molecule dyes suitable for single-molecule imaging (Figure 1D; (Grimm et al., 2015)). To examine the effect of tagging CTCF and Rad21, which are both essential proteins, we performed control experiments in the doubly-tagged mESC line (C59), and observe no effect on mESC pluripotency (Figure S1), expression of key stem cell genes (Figure S2A) or tagged protein abundance (Figure S2B). Next, we performed chromatin immunoprecipitation followed by DNA sequencing (ChIP-Seq) using antibodies against CTCF and Rad21 in both wild-type (wt) and the double knock-in C59 line. To further validate our endogenous tagging approach, we compared ChIP-Seq enrichment for both wt and C59 at called wt peaks and observe similar enrichment (Figure 1E-F). Notably, 97% of called Rad21 peaks co-localize with a called CTCF peak (Figure S3–4; table S1), largely confirming previous reports of ~70% overlap (Parelho et al., 2008; Wendt et al., 2008). However, chromatin co-occupancy by ChIP-seq at the same sites does not necessarily mean that CTCF and Rad21 bind simultaneously. Thus, to determine whether CTCF and cohesin physically interact, we performed co-immunoprecipitation (co-IP) studies. CTCF IP pulled down cohesin subunits Rad21, Smc1 and Smc3 in both wt and C59 mES cells (Figure 1G), demonstrating a physical interaction between CTCF and cohesin, which is not affected by endogenous tagging. Taken together with the previous literature (Dekker and Mirny, 2016; Dixon et al., 2012; Flavahan et al., 2015; Ghirlando and Felsenfeld, 2016; Hnisz et al., 2016a; Lupianez et al., 2015; Merkenschlager and Nora, 2016; Nora et al., 2012; Rao et al., 2014; Sanborn et al., 2015; Skibbens, 2016; Wang et al., 2016; de Wit et al., 2015) implicating CTCF and cohesin in chromatin looping, we propose that CTCF and cohesin form a “Loop Maintenance Complex” (LMC; Figure 1H).

To investigate the dynamics of the LMC we measured the residence time of CTCF and cohesin on chromatin. First, we used highly inclined and laminated optical sheet illumination (Tokunaga et al., 2008) (Figure 2A) and single molecule tracking (SMT) to follow single Halo-CTCF molecules in live cells. By using long exposure times (500 ms), to “motion-blur” fast moving molecules into the background (Chen et al., 2014), we could visualize and track individual stable CTCF binding events (Figure 2B; Movie S1). We recorded thousands of binding event trajectories and calculated their survival probability. A double-exponential function, corresponding to specific and non-specific DNA binding (Chen et al., 2014), was necessary to fit the Halo-CTCF survival curve (Figure 2C). After correcting for photo-bleaching (Figure S5A), we estimated an average residence time (RT) of ~1 min for CTCF in both mES and U2OS cells (Figure 2D). DNA-binding defective CTCF mutants or Halo-NLS alone interacted very transiently with chromatin (RT ~1 s; Figure 2D). The measured RT did not depend on the dye or exposure times (Figure S5B). We note that a CTCF RT of ~1 min is a genomic average and that some binding sites likely exhibit a slightly longer or shorter mean residence time. To cross-validate these results using an orthogonal technique, we performed fluorescence recovery after photo-bleaching (FRAP) on Halo-CTCF and quantified the dynamics of recovery (Figure S6A). Both Halo-CTCF in mES cells (Figure 2E) and Halo-hCTCF in U2OS cells (Figure S6C) exhibited FRAP recoveries consistent with a RT of ~1 min. Our results differ considerably from a previous CTCF FRAP study using over-expressed transgenes, which reported rapid 80% recovery in 20 s (Nakahashi et al., 2013). However, when we used similar transiently over-expressed Halo-CTCF instead of endogenous knock-in cells, we also observed similarly rapid recovery (Figure S6B), suggesting that over-expression of target proteins can result in artefactual measurements. This finding underscores the importance of studying endogenously tagged and functional proteins. Thus, although CTCF (RT~1 min) binds chromatin much more stably than most sequence specific transcription factors (RT~2–15 s) (Chen et al., 2014; Mazza et al., 2012), its binding is still highly dynamic.

**Figure 2.**
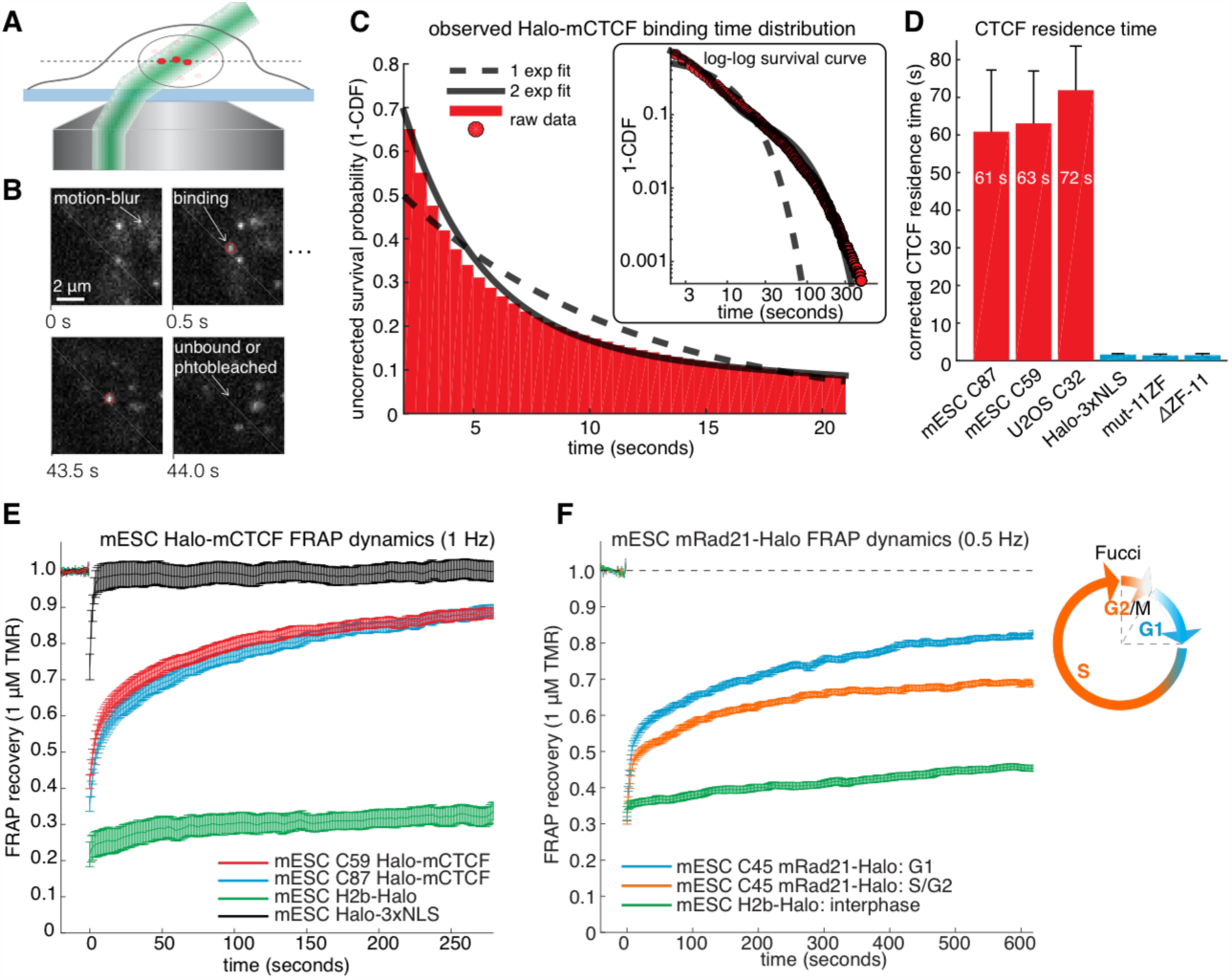
CTCF and Cohesin have very different residence times on chromatin. **(A)** Sketch illustrating HiLo illumination. **(B)** Example images showing single Halo-CTCF molecules labeled with JF549 binding chromatin in a live mES cell. **(C)** A plot of the uncorrected survival probability of single Halo-CTCF molecules and one-and two-exponential fits. Right inset: a log-log survival curve. **(D)** Photobleaching-corrected residence times for Halo-CTCF, Halo-3xNLS and a zinc-finger (11 His→Arg point-mutations) mutant or entire deletion of the zinc-finger domain. Error bars show standard deviation between replicates. For each replicate we recorded movies from 4–8 cells and calculated the average residence time using H2b-Halo for photobleaching correction. Each movie lasted 20 min with continuous low-intensity 561 nm excitation and 500 ms camera integration time. Cell were labeled with 1–100 pM JF549. **(E)** FRAP recovery curves for Halo-mCTCF, H2b-Halo and Halo-3xNLS in mES cells labeled with 1 μM Halo-TMR. **(F)** FRAP recovery curves for mRad21-Halo and H2b-Halo in mES cells labeled with 1 μM Halo-TMR. Right: sketch of Fucci cell-cycle phase reporter. Each FRAP curve shows mean recovery from >15 cells from ≥3 replicates and errorbars show the standard error.

We next investigated the cell-cycle dependent cohesin binding dynamics (Gerlich et al., 2006). In addition to its suspected role in holding together chromatin loops, cohesin mediates sister chromatid cohesion from replication in S-phase to mitosis. Thus, since TAD demarcation is strongest in G1 before S-phase (Naumova et al., 2013), we reasoned that cohesin dynamics in G1 should predominantly reflect the chromatin looping function of cohesin. To control for the cell-cycle, we deployed the Fucci system to distinguish G1 from S/G2-phase using fluorescent reporters (Figure S7; Supplementary Materials) in the C45 and C59 mESC lines. We then performed FRAP on mRad21-Halo (Figure 2F) and mRad21-SNAP_f_ (Figure S6E). We observed much slower recovery of mRad21 compared to CTCF, but nevertheless a significantly faster recovery in G1 than in S/G2-phase. The slow Rad21 turnover precluded SMT experiments (Supplementary Materials). Model-fitting of the G1 mRad21 FRAP curves (Figure S6F-G) revealed an RT~22 min. Previous cohesin FRAP studies have reported RTs that differ by two orders of magnitude (Supplementary Materials): as was seen for CTCF, over-expressed mRad21-Halo also showed much faster recovery than endogenous Rad21-Halo (Figure S6H). Although kinetic modeling of FRAP curves should be interpreted with some caution (Mazza et al., 2012), these results nevertheless demonstrate a surprisingly large (~20x) difference in RTs between CTCF and cohesin, which is difficult to reconcile with the notion of a biochemically stable LMC assembled on chromatin. However, it is still possible that CTCF and cohesin form a stable complex in solution when not bound to DNA.

To investigate this possibility, we analyzed how CTCF and cohesin each explore the nucleus. Tracking fast-diffusing molecules has been a major challenge. To overcome this issue, we took advantage of bright new dyes (Grimm et al., 2016) and developed stroboscopic (Elf et al., 2007) photo-activation single-molecule tracking (paSMT; Figure S8A), which makes tracking unambiguous (Supplementary Materials). We tracked individual Halo-CTCF molecules at ~225 Hz and plotted the displacement between frames (Figure 3A). Most Halo-CTCF molecules exhibited displacements similar to our localization error (~35 nm; Supplementary Materials) indicating chromatin association, whereas a DNA-binding defective CTCF mutant exhibited primarily long displacements consistent with free diffusion (Figure 3B; Movie S2–3). To characterize the nuclear search mechanism, we performed kinetic modeling of the measured displacements (Figure S8B; Supplementary Materials; (Mazza et al., 2012)) and found that in mES cells, ~50% of CTCF is bound to cognate sites, ~19% is non-specifically associated with chromatin (e.g. sliding or hopping) and ~31% is in free 3D diffusion (table S2). Thus, after dissociation from a cognate site, CTCF searches for ~65 s on average before binding the next cognate site: ~65% of the total nuclear search is random 3D diffusion (~41 s on average), whereas ~35% (~24 s on average) consists of intermittent non-specific chromatin association (e.g. 1D sliding; table S2). The nuclear search mechanism of CTCF in human U2OS cells was similar albeit slightly less efficient (table S2; Figure S8F). We note that CTCF’s search mechanism, with similar amounts of 3D diffusion and 1D sliding, is close to optimal according to the theory of facilitated diffusion (Mirny et al., 2009).

**Figure 3.**
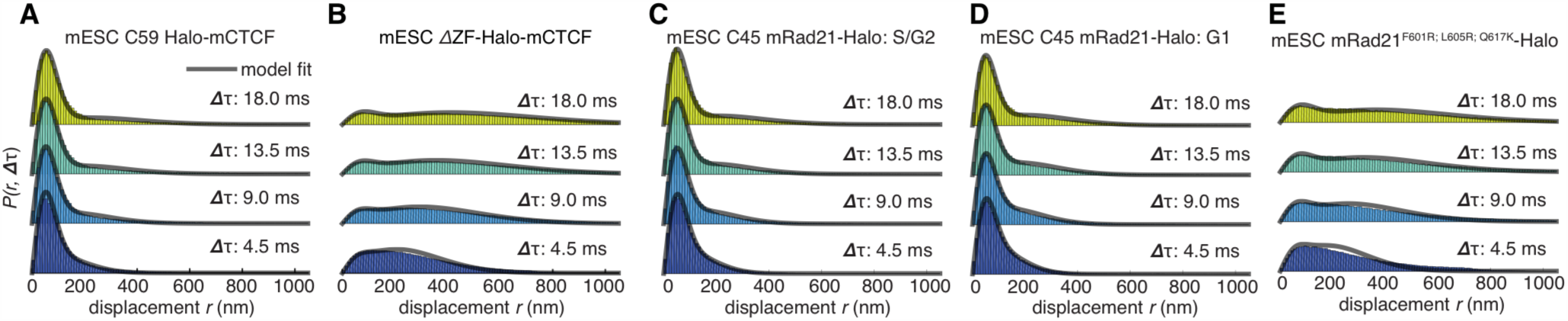
Dynamics of CTCF and cohesin’s nuclear search mechanism. Top row: single-molecule displacements from 225 Hz stroboscopic (single 1 ms 633 nm laser pulse per camera integration event) paSMT experiments over multiple time scales for (**A**) C59 Halo-mCTCF, (**B**) a Halo-mCTCF mutant with the zinc-finger domain deleted, C45 mRad21-Halo in S/G2 phase (**C**) and G1 phase (**D**) and (**E**) a Rad21 mutant that cannot form cohesin complexes. Kinetic model fits (3 fitted parameters) to raw displacement histograms are shown as black lines. All calculated and fitted parameters are listed in table S2.

Similar analysis of mRad21-Halo in G1 and S/G2 (Figure 3C-D) revealed that cohesin complexes diffuse rather slowly compared to CTCF (table S2) and that roughly half of cohesins are topologically engaged with chromatin (G1: ~40%; S/G2: ~50%) compared to ~13% in non-specific chromatin association and the remainder in 3D diffusion (G1: ~47%; S/G2: ~57%). Conversely, a Rad21 mutant (Haering et al., 2004) unable to form cohesin complexes displayed rapid diffusion and little chromatin association (Figure 3E; Movie S4–5). Like this Rad21 mutant, overexpressed wild-type mRad21-Halo also showed negligible chromatin association (Figure S8E) again underscoring the importance of studying endogenously tagged proteins at physiological concentrations. Topological association and dissociation of cohesin is regulated by a complex interplay of co-factors such as Sororin and Wapl (Skibbens, 2016). If we nevertheless apply a simple two-state model to analyze cohesin dynamics (Supplementary Materials), we estimate an average search time of ~33 min in between topological engagements of chromatin in G1, with ~77% of the search time spent in 3D diffusion (~26 min) compared to ~23% in non-specific chromatin association (7 min). Thus, for each specific topological cohesin chromatin binding-unbinding cycle in G1, CTCF binds and unbinds its cognate sites ~25–30 times. These results are certainly not consistent with a model wherein CTCF and cohesin form a stable LMC. Moreover, since CTCF diffuses much faster than cohesin (table S2), it also seems unlikely that CTCF and cohesin form stable complexes in solution.

To resolve these apparently paradoxical findings, we investigated the nuclear organization of CTCF and cohesin simultaneously in the same nucleus. We labeled Halo-mCTCF and mRad21-SNAP_f_ in C59 mES cells with the spectrally distinct dyes JF646 and JF549 (Grimm et al., 2015), respectively, and performed two-color direct stochastic optical reconstruction microscopy (dSTORM) super-resolution imaging in paraformaldehyde-fixed cells (Figure 4A). We localized individual CTCF and Rad21 molecules with a precision of ~20 nm, less than half the size of the cohesin ring. We observe significant clustering of both CTCF and Rad21 (Figure 4A and S9A-C). We next confirmed clustering using photo-activation localization microscopy (PALM) and found that both CTCF and Rad21 predominantly form small clusters (Fig 4B and S9; mean cluster radius ~30–40 nm). To determine whether individual CTCF and cohesin molecules co-localize, we calculated the pair cross correlation, *C*(*r*) (Stone and Veatch, 2015). *C*(*r*) quantifies spatial co-dependence as a function of length, *r*, and *C*(*r*) = 1 for all *r* under complete spatial randomness (CSR). CTCF and cohesin exhibited significant co-localization (*C*(*r*)>1) at very short distances in mES cells (Figure 4C). Conversely, CTCF and cohesin were nearly independent at length scales beyond the diffraction limit, emphasizing the importance of super-resolution approaches. A mES cell line co-expressing H2b-SNAP_f_ and Halo proteins imaged under the same dSTORM conditions showed no cross-correlation (Figure 4C), thereby ruling out technical artifacts. Thus, our two-color dSTORM results provide compelling evidence that a large fraction of CTCF and cohesin indeed co-localize at the single-molecule level consistent with the LMC model and reveals a clustered nuclear organization.

**Figure 4.**
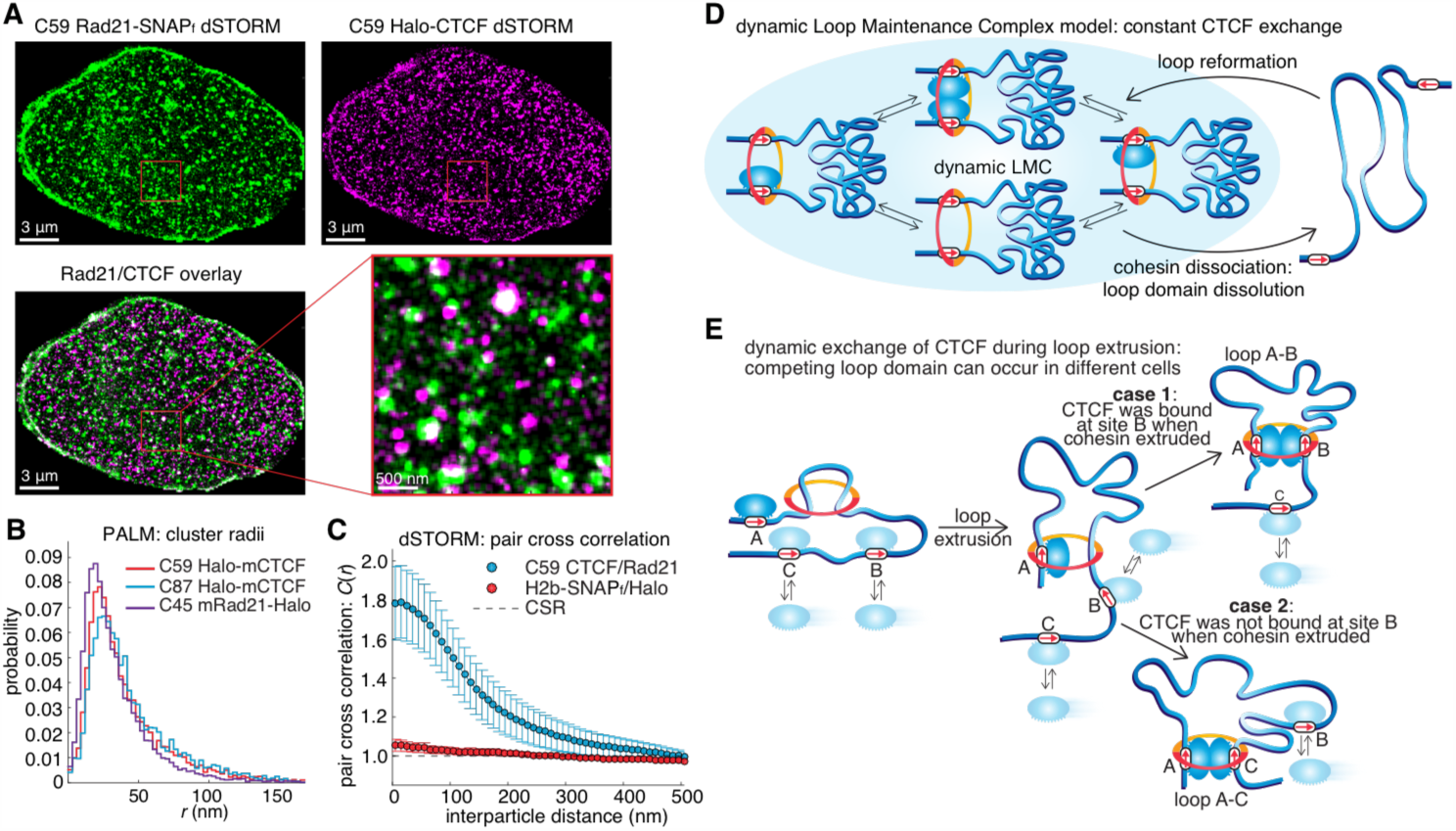
Models of CTCF/cohesin mediated chromatin loop dynamics. **(A)** Two-color dSTORM of C59 mESCs with mRad21-SNAP_f_ labeled with 500 nM JF549 (green) and Halo-mCTCF labeled with 500 nM JF646 (magenta). High-intensity co-localization is shown as white. Low intensity co-localization is not visible. Zoom-in on red 3 μm square. **(B)** Cluster radii distributions for CTCF (C87 and C59) and Rad21 (C45) from single-color PALM experiments. **(C)** Pair cross correlation of C59 and mESC H2b-SNAP_f_ co-expressing Halo-only. Errorbars are standard error from 12–18 cells over three replicates. **(D)** Sketch illustrating the concept of a dynamic Loop Maintenance Complex (LMC) composed of CTCF and cohesin with constant CTCF exchanging and slow, rare cohesin dissociation, which causes loop deformation and topological re-orientation of chromatin. **(E)** Sketch illustrating how dynamic CTCF exchange during loop extrusion of cohesin may explain alternative loop formations when two competing convergent sites (B and C) for another site (A) exist.

## Discussion

Chromatin loop domains are widely believed to be very stable structures (Andrey et al., 2016; Ghirlando and Felsenfeld, 2016; Hnisz et al., 2016b) permanently held together by a LMC composed of two CTCFs and cohesin. While our *in vitro* biochemical (Figure 1G) and co-localization (Figure 4A-C) experiments do demonstrate complex formation between CTCF and cohesin, our SMT experiments paradoxically reveal this complex to be highly transient and dynamic (Figure 2-3). To reconcile these observations, we therefore propose a “dynamic LMC” model where CTCF mainly functions to position cohesin at loop boundaries, whereas cohesin physically holds together the two chromatin strands. While cohesin holds together a given chromatin loop, different CTCF molecules are frequently alighting and departing in a dynamic exchange thus giving rise to a “transient protein complex” with a molecular stoichiometry that cycles over time (Figure 4D). Since topological association of cohesin is infrequent (~33 min in G1), dissociation of cohesin (~22 min) likely causes the loop to fall apart (Figure 4D). Even if larger clusters of CTCF and cohesin (Figure 4A-C; Figure S9) hold together loop domains, their lifetimes are unlikely to be more than 1–2 hours. Thus, our results suggest that chromatin loops are continuously formed and dissolved throughout the cell cycle. This dynamic view of loops is also more consistent with polymer simulation studies, which imply that static loop domains cannot reproduce experimentally observed chromatin interaction frequencies (Fudenberg et al., 2016; Sanborn et al., 2015). We note that our quantitative characterization of CTCF and cohesin dynamics could be useful for parameterizing future polymer models. While our results reveal loops to be highly dynamic, the question of how they are formed remains. An attractive but unverified recent model suggests that loops are formed by cohesin-mediated loop extrusion (Fudenberg et al., 2016; Sanborn et al., 2015), whereby cohesin extrudes a loop by sliding on DNA (Davidson et al., 2016; Stigler et al., 2016) until it encounters two convergent and bound CTCF sites (Figure 4E). In the context of the loop extrusion model, our results suggest that dynamic and stochastic CTCF occupancy at cognate CTCF sites may explain the formation of competing loop domains (Figure 4E). This would also explain why DNA-FISH measurements show that most loops are only present in a subset of cells at any given time (Sanborn et al., 2015; Williamson et al., 2014). Finally, the highly dynamic view of frequently breaking and forming chromatin loops presented here may also facilitate dynamic long-distance enhancer-promoter scanning of DNA in *cis,* which may be critical for temporally efficient regulation of gene expression.

## Acknowledgements

We thank Luke Lavis for generously providing JF dyes, Gina M. Dailey for extensive assistance with cloning, Astou Tangara for microscopy assembly and maintenance and Kartoosh Heydari for flow cytometry assistance. We also thank members of the Tjian and Darzacq labs, Douglas Koshland and Miriam Huntley for insightful comments on the manuscript. This work was performed in part at the CRL Molecular Imaging Center, supported by the Gordon and Betty Moore Foundation. This work used the Vincent J. Coates Genomics Sequencing Laboratory at UC Berkeley, supported by NIH S10 Instrumentation Grants 10RR029668 and S10RR027303. ASH is a postdoctoral fellow of the Siebel Stem Cell Institute. This work was supported by NIH grants UO1-EB021236 and U54-DK107980 (XD), the California Institute of Regenerative Medicine grant LA1-08013 (XD and RT), and by the Howard Hughes Medical Institute (003061, RT). ChIP-Seq data has been deposited at NCBI GEO.

